# How the human brain introspects about one’s own episodes of cognitive control

**DOI:** 10.1101/208637

**Authors:** David Soto, Mona Theodoraki, Pedro M. Paz-Alonso

**Affiliations:** Basque Center on Cognition, Brain and Language, San Sebastian, Spain.; Ikerbasque, Basque Foundation for Science, Bilbao, Spain.; Division of Brain Sciences, Imperial College London, United Kingdom

**Author notes:** Correspondence to, Basque Center on Cognition, Brain and Language, Paseo Mikeletegi 69, 2nd Floor 20009 San Sebastian.

## Abstract

Metacognition refers to our capacity to reflect upon our experiences, thoughts and actions. Metacognition processes are linked to cognitive control functions that allow keeping our actions on-task. But it is unclear how the human brain builds an internal model of one’s cognition and behaviour. We conducted 2 fMRI experiments in which brain activity was recorded ‘online’ as participants engaged in a memory-guided search task and then later ‘offline’ when participants introspected about their prior experience and cognitive states during performance. In Experiment 1 the memory cues were task-relevant while in Experiment 2 they were irrelevant. Across Experiments, the patterns of brain activity, including frontoparietal regions, were similar during on-task and introspection states. However the connectivity profile amongst frontoparietal areas was distint during introspection and modulated by the relevance of the memory cues. Introspection was also characterized by increased temporal correlation between the default-mode network (DMN), frontoparietal and dorsal attention networks and visual cortex. We suggest that memories of one’s own experience during task performance are encoded in large-scale patterns of brain activity and that coupling between DMN and frontoparietal control networks may be crucial to build an internal model of one’s behavioural performance.

## 1 Introduction

A fundamental constituent of the human condition is the ability to reflect upon our own ongoing or recently past cognition and behavioural states. This ability is known as metacognition. For instance, metacognitive accounts assume that the brain builds and maintains an internal model of one’s own cognition and behaviour and that experience-dependent updates of this model (e.g. following a performance error) may have consequences for behavioural control (Nelson & Narens, 1990). Processes of cognitive control are also linked to metacognition in that control allows us to keep our thoughts and actions aligned with the task at hand. Both self-reflection and control are fundamental to regulate behaviour. Self-monitoring may change the behaviour of one’s cognitive system and promote adaptive behaviour and learning (Nelson & Narens, 1990). However, current understanding of the relationship between metacognition, and cognitive control functions is far from complete. Here we used functional MRI to assess the neural correlates of introspection about one’s own recently past episodes of attentional control during task performance. Using a visual search paradigm, we ask to what extent the brain reassembles one’s own previous cognitive control states via introspection, namely, how the prior on-task states during search are re-experienced offline when participants are no longer engaged in the task and in the absence of the same external conditions.

The focus of this work is therefore one component of metacognition that relates to the ability to introspect about one’s own ongoing, or recently past, cognitive states and behavioural processes. It has been argued that introspection is exclusively related to conscious states(Overgaard & Sandberg, 2012; Soto & Silvanto, 2014) while at least some other metacognitive components that monitor ongoing behaviour may operate in a more automatic manner and even independently to some extent of conscious awareness (Charles et al., 2017; Jachs, Blanco, Grantham-Hill, & Soto, 2015; Kanai, Walsh, & Tseng, 2010; Shea et al., 2014). Introspection is here used to refer to the conscious observation of one’s own cognition and behaviour, rather than memories that we have about the states of the world. This view is in keeping with theories that emphasize the critical importance for conscious experience of those superordinate mental states that represent oneself as being in the relevant first-order mental state (e.g. of performing a task; cf. (Rosenthal, 1986). Hence, the term introspection here refers to the non-automatic, effortful process that brings knowledge about one’s own recently past mental processes and events (Schwitzgebel, 2016), which in the context of the present study will be the individual experience of exerting cognitive control following a series of task events.

The internal model of one’s own cognition and behaviour that is the object of introspection requires the integration of information from multiple brain systems (e.g. perception, attention, decision and action). Hence, in keeping with a network memory approach (Fuster, 1997, 2009) that proposes that memory is a property of all neural systems reflected in global patterns of brain activity and connectivity for a given experience and behaviour, it is here hypothesized that introspection about one’s own recently past episodes of attentional control may be mediated by the re-enactment of similar large-scale patterns of brain activity engaged during the actual behavioural performance. Presumably, therefore, introspection about one’s own recently past episodes of attentional control may be mediated by re-instatement of activity in dorsal and ventral frontoparietal networks that are involved in external attention (Corbetta & Shulman, 2002). This is our first hypothesis.

The default-mode network (DMN) is thought to be involved in different types of self-reflective or internally directed processes, including self-related remembering and thinking about the oneself in a future context (Gusnard, Akbudak, Shulman, & Raichle, 2001). Self-related processing may be assumed to operate during ‘rest’ periods and hence attenuate during task performance. This is further suggested by reductions in neural activity in default-mode regions during task engagement, alongside with response increases in task ‘positive’ networks such as the frontoparietal control network and the dorsal attention network. Indeed, anticorrelation between key nodes of the DMN (e.g. posterior cingulate, medial prefrontal cortex) and dorsal attention networks has been assumed to reflect an intrinsic property brain organisation to support opposing functions (e.g. external attention vs. reflection). However, recent evidence suggests this view may be too simplistic. Although the DMN is involved in a variety of internally-directed processes (Gusnard et al., 2001), different studies demonstrate co-activation and positive functional connectivity between the DMN and the frontoparietal network during different task conditions including working memory, memory retrieval, reasoning, theory of mind and attention (Christoff, Gordon, Smallwood, Smith, & Schooler, 2009; Fox, Spreng, Ellamil, Andrews-Hanna, & Christoff, 2015; Dixon et al., 2017; Legrand & Ruby, 2009; Crittenden, Mitchell, & Duncan, 2015; Leech, Kamourieh, Beckmann, & Sharp, 2011). Interestingly, Christoff and colleagues (Christoff, Cosmelli, Legrand, & Thompson, 2011) also proposed the idea that self-referential processes relate not only to the personal attributes of oneself but also to the perception of oneself in a sensory-motor context (e.g. the self as an agent that interacts with the world). An interesting possibility therefore that will be tested here is that functional coupling between the DMN and ‘task-positive’ control networks is enhanced during introspection of one’s own recent episodes of attention control during task performance.

## 2 Methods

### 2.1 Participants

Nineteen undergraduate students were recruited (mean age of 22 years; nine female). All participants gave informed written consent to take part in accordance with the terms of approval granted by the local research ethics committee and the principles expressed in the Declaration of Helsinki, and were naive to the purpose of the experiment. All participants reported normal or corrected-to-normal vision, no history of neurological disease, and no other contraindications for MRI

### 2.2 Experimental task and procedure

Participants came to the laboratory 1-2 days prior to scanning to receive information about the study, give informed consent and receive training in the task. Each trial started with a fixation cross for 0.5 seconds, followed by a 0.5 s blank period. There followed the presentation of one of eight possible Japanese hiragana symbols (the cues) for 0.5 s. Then, a blank screen was presented for 1.5 s delay, which was followed by a search display comprising four outline circles in red, green, yellow and blue displayed at the corners of an imaginary square. Three of the coloured circles contained a vertical line and the remaining circle contained a tilted line (the search target). The search task was to discriminate whether the target line was oriented to the left or to the right from the vertical. Search display appeared for 0.1 s to discourage eye movements. Participants were instructed that 4 of those hiragana cues were predictive about the color of the circle surrounding the target (the valid condition), while the remaining 4 were not associated with any relevant color. The four predictive hiragana cues were randomly selected for each participant. The same hiragana cues were used in the training phase outside the scanner and also during the scanning session. Participants were encouraged to learn the association between each of the relevant hiragana cues and the colours in the search array in order to improve their search performance. Participants learned these contingencies prior to the scanning session and by the end of the training protocol, search performance was faster on valid relative to neutral trials (data not shown). The experimental task is shown in Figure 1.

FMRI Experiment 1 (see Figure 1A) started with the search task in which four of the hiragana cues were predictive of the colour of the circle surrounding the search target while the remaining 4 hiragana cues were neutral and not associated with any target feature. The colour of the circle surrounding the search target was randomly selected on these non-predictive trials. A central fixation point was presented during the inter-trial interval. Following the search task, participants were required to introspect. Here brain responses were recorded in the absence of any over task requirement, when observers introspected about prior search behaviour in the context of each of the hiragana cues. In this second scanning run, following the presentation of each hiragana cue, participants were asked to re-experience in their mind the prior cognitive and behavioural states associated with performance for that particular hiragana cue. These introspection trials presented observers with an hiragana cue followed by a blank screen while keeping the trial timings of the overt search task blocks.

The trials of the task within each scanning run were organised in mini-blocks of 4 trials each, either of Cued (C) or neutral (N) trials, with block order counterbalanced as follows: C-N-C-N-N-C-C-N-N-C-C-N-N-C-N-P. Hence there were a total of 64 trials per run (32 per validity condition). Between each mini-block there appeared a instruction display for 1.5 seconds indicating the validity status of the cue in the next mini-block, which was followed by a further 4.5 s delay. A block design was used to separate on-task and introspection states. The first scanning run was on-task and the second scanning run was introspection (see Figure 1).

**Figure 1:**
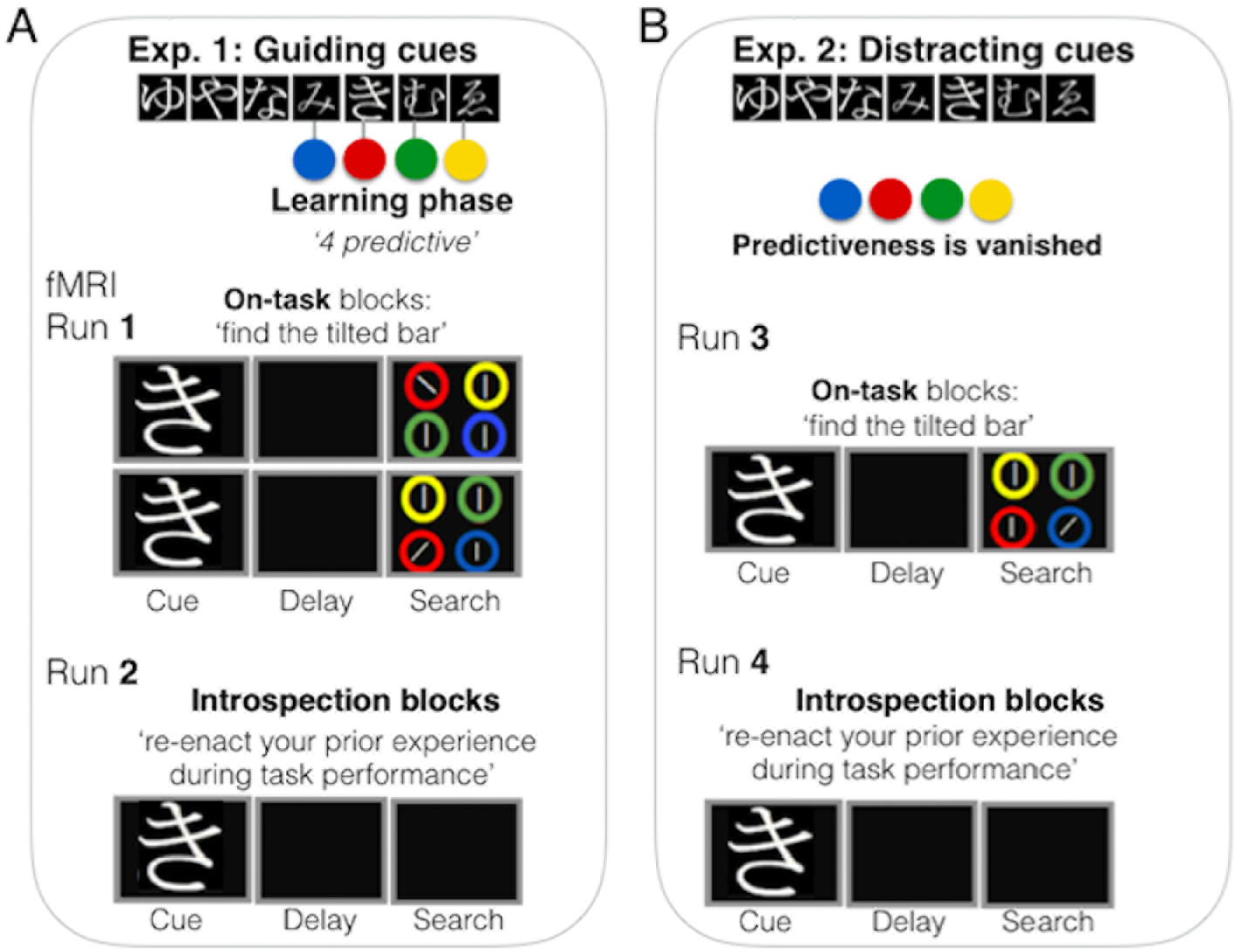
Illustration of the experimental protocol in Experiment 1 (A) and Experiment 2 (B).

We elected to use abstract hiragana cues to ensure that neural responses during introspection of the previous trials on-task relied on the relevance or attentional validity of the cue for the search task (i.e. an attention episode comprised of associative links between an abstract cue and a specific feature and states of control during search), rather than on re-activation or priming of low-level sensory processes associated with specific feature cues. The use of abstract hiragana cues was also crucial in experiment 2 when the validity of the hiragana cues was extinguished (see below).

In Experiment 2, the validity of the hiragana cues for the search task was extinguished and accordingly the colour previously associated with a given hiragana cue now matched a distracter in search (the invalid condition). There were four additional neutral cues that had no association with the colour matching the search target during this phase. Participants were informed prior to the beginning of Experiment 2 that the predictiveness of the cues vanished and that they were no longer associatted with the colour surrounding the tilted target. The structure of the mini-blocks was the same as before, except for the change in the contingencies noted above; also between mini-blocks there appeared a instruction remainder that the cues were no longer predictive. The first scanning run of Experiment 2 comprised on-task trials. This was followed by a second run of introspection which had the same structure except that the search displays were absent. Here, following the presentation of each cue, participants here were required to introspect about their experience and cognitive states associated with the task performance in this context of invalid cueing.

Both Experiment 1 and 2 were carried out in the same scanning session.

Upon completion of the scanning protocol, participants were asked to report about their experience inside the scanner during the introspection blocks. Overall, participants’ reports were highly consistent. They reported to have experienced the visual scene with the coloured circles and lines, remembered how the hiragana cues triggered color associations, and how these were used to explore the display, find the search target and respond to it.

### 2.3 Functional MRI protocol

Functional volumes consisted of multi-slice T2^*^-weighted echoplanar images (EPI) with blood oxygenation level dependent (BOLD) contrast with multiband acceleration factor of 4. We used the following scanning parameters to achieve whole brain coverage: TR = 1000 ms, TE = 54.8 ms, 40 coronal slices, 3.2 mm slice thickness, 0 interslice gap, and FoV = 205 × 205 mm. To facilitate anatomical localization and cross-participant alignment, a high-resolution whole-brain structural T1-weighted, magnetization prepared rapid gradient echo (MP-RAGE) scan was acquired for each participant

### 2.4 Functional MRI Data Analyses: GLM

We used FEAT (fMRI Expert Analysis Tool) Version 6.0, as part of FSL (www.fmrib.ox.ac.uk/fsl). The first 10 EPI volumes were removed to account for T1 equilibrium effects. Non-brain removal was performed using Brain Extraction Tool. Motion correction of functional scans was carried out using FM-RIB’s Linear Image Registration Tool MCFLIRT. We applied a 100 s high-pass temporal filtering to remove low frequency noise, and spatial smoothing using a FWHM Gaussian kernel of 5 mm. Time-series statistical analyses were conducted using FILM (FMRIB’s Improved Linear Model) with local autocorrelation correction. The data were analyzed using voxelwise time series analysis within the framework of the general linear model. A design matrix was generated with a double-gamma hemodynamic response function and its first temporal derivative. Trials were modelled from the onset of the hiragana symbols with duration of 2.1 s (including 0.5 s cue exposure, 1.5 s blank screen delay and 0.1 s search display. We separately modelled the predictive and non-predictive cues. The 1.5 s instruction prior to each of the task and introspection blocks and the presence of catch trials and error responses were modelled as regressors of no interest. Standard motion parameters were also included in each individual subject’s general linear model as further regressors of no interest.

We obtained contrasts of parameter estimates for cued (i.e. valid in Experiment 1, invalid in Experiment 2) > (<) the neutral condition for each individual. Data from the on-task condition was carried forward to a higher-level analysis using FLAME 1+2 (FMRIB’s Local Analysis of Mixed Effects) in the form of one-sample t-test to define consistent brain regions of activation across participants. The data were registered to individual high-resolution structural images using FLIRT Boundary-Based Registration, and then co-registered into standard MNI space for visualization. Clusters of activity in the on-task condition were identified by a voxelwise threshold of Z = 2.3 and a cluster significance threshold of P=.05, corrected for multiple comparisons (using Gaussian Random Field Theory).

### 2.5 Pattern similarity analyses

To assess the similarity of brain activity patterns during introspection and on-task states we first used the unthresholded second-level group-average contrast of parameter estimates (cued>neutral) associated with the task and introspection effects and used Pearson correlation to compute the similarity of voxel estimates across the spatial extent occupied by the relevant clusters. This was done first using the total brain area activated by the task as a whole-brain mask and also using masks for each of the clusters of activity identified in the on-task state. This approach provides a similarity metric of the group-averaged activation maps. In order to confirm the pattern of results, we also performed a second analyses that accounted for inter-subject variance. To this end, we calculated the Pearson correlation between on-task and introspection space separately within each subject. Since the correlation coefficient is inherently restricted to a range from −1 to +1, an arc-hyperbolic tangent transform (Fisher, 1921) was applied. Then, a one-sample t-test was carried out on these z-scores to test the statistical significance of the similarity estimates.

### 2.6 Beta series correlations

Firstly, we assessed functional connectivity in fronto-parietal areas via the beta-series correlation method (Rissman et al., 2004), implemented in SPM8 with custom Matlab scripts. Beta series correlation analyses is an established method which presents a series of advantages that are relevant for our study, compared to other voxel-wise analyses of connectivity such as psychophysiological interactions (PPI). These include 1.- The beta-series approach do not present the constrain of the PPI regressor being highly correlated to psychological task regressor, which might reduce the power to detect effects of interest when using PPI. 2.- Beta series is a method specifically designed for slow event-related designs, which is akin to the present design in which the different conditions were presented in a set of mini-blocks lasting for around 14.5 seconds; 3.- Beta series is a flexible method that capitalizes on a GLM applied to beta images obtained for each single event/epoch that allows to examine seed-to-voxel and ROI-to-ROI functional connectivity as a function of the different task variables. This is a clear advantage relative to to PPI, whichÂă can only model contributions of a single area at a time. Here we elected to use the beta-series method because it was specially suitable for our research questions and appropriate for our fMRI experimental design.

For the beta-series functional connectivity analyses we used restrictive motion correction procedures typically used in SPM. First, ArtRepair (Center for Interdisciplinary Brain Sciences Research, Stanford Medicine; Mazaika et al., 2009) was used to correct scan-to-scan motion >=0.5 mm and scan-to-scan variation in global intensity >= 1.3 %. Importantly, none of the subjects had more than 20 % of to-be-repaired scans across functional runs. Second, to control for any residual linear motion effects, the 6 motion regressors (i.e., x, y, z, pitch, yaw and roll) were included in the GLM used for functional connectivity analysis as covariates of non-interest (additional details on the motion correction pipeline can be found in the Supplemental Materials).

Pairwise functional connectivity between frontoparietal regions was examined. To do so, 5-mm-radius spheres were build centered at the highest local maxima observed in the t-contrast All-Null across all subjects, q < .001 FDR voxel-wise corrected for multiple comparisons. Note that for the functional connectivity analyses we elected to use the t-contrast All-Null in order to avoid biases in connectivity that could arise if the ROIs were derived from the cuing contrast (i.e. cued><uncued) contrast. Note that the pattern similarity analyses assessed the extent to which the cueing effect parameter estimates on-task resembled those in the introspection period, hence the need of using this conttast for the similarity analyses.

The ROIs were located in left inferior parietal cortex (IPC; −39 −46 52; BA 40; T-value = 7.27), left superior parietal cortex (SPC; −18 −58 49; BA 7; T-value = 6.16), left middle frontal gyrus (MFG; −54 11 37; BA 9; T-value = 8.23), right IPC (51 −37 46; BA 40; T-value = 8.55), right SPC (24 −67 55; BA 7; T-value = 5.43), and right MFG (57 11 34; BA 9; T-value = 9.43). The relative superior/ inferior loci of the parietal ROIs was determined according to the Automated Anatomical Labeling (AAL) available in SPM.

Beta-series correlation values were calculated for each pair of ROIs, participant and experimental condition. Then, Fisher’s z-score transformed beta-series correlation values between left frontoparietal regions (i.e., IPC, SPC, MFG) and between right frontoparietal regions (i.e., IPC, SPC, MFG) were averaged and submitted to repeated-measures analyses of variance (ANOVAs) including Hemisphere (left, right), condition (task, introspection) and cueing (cue, neutral) as within-subjects independent measures, and averaged functional connectivity among nodes as the dependent measure. Two identical ANOVAs were conducted separately for Experiment 1 and 2.

### 2.7 Temporal correlation analyses between brain networks

An analysis of the inter-relations among large-scale brain networks was carried out following a dual regression approach implemented in FSL (see below). Smith and colleagues (Smith et al., 2009) identified several brain networks using independent component analyses of a large sample, which were given functional labels based on their correspondence to the BrainMap database of human fMRI studies. These include the default mode network, 3 visual networks (medial, occipital pole and lateral), a cerebellum network, auditory, sensorimotor, executive control network, and left and right frontoparietal networks. Also, two more networks from Smith et al. (2009), that correspond to the dorsal attention networks (i.e. DAN1 and DAN2) were included in a dual regression approach (12 networks in total) to study between network functional connectivity in our dataset. ICA-AROMA (Pruim et al., 2015) was used for removing components associated with motion artifacts from each individual fMRI dataset, which could contaminate the analyses of the temporal correlations between brain networks. ICA-AROMA was carried following realigment of the scans with MCFLIRT, and smoothing, and the dual regression method was applied on this output following high-pass filtering (Pruim et al., 2015). In this dual regression approach, the time courses of seach subject’s network, and, for each on-task and introspection conditions were extracted by back-projecting the canonical networks from Smith et al. (2009) into each individual fMRI dataset using a general linear model (Beckmann, Mackay, Filippini, & Smith, 2009). This involves using the spatial maps of the canonical networks as regressors, and then using the derived time courses to define subject specific maps associated with the canonical networks. Then, the correlation between the time course of the DMN and the rest of the networks was computed using Pearson correlation for each subject, both during on-task and introspection periods. T-tests were carried of Fisher z-scored transformed correlations to assess consistent changes in network dynamics across subjects as a function of the experimental context (i.e. on-task vs introspection) (see (Roseman, Leech, Feilding, Nutt, & Carhart-Harris, 2014) for a similar approach. We also note the correlations that survive FDR correction for multiple tests (Benjamini & Hochberg, 1995).

## 3 Results

### 3.1 Search Performance

Median search latencies of the correct responses in Experiment 1 were faster on valid (mean = 537.2, SEM = 12.37) relative to neutral trials (mean = 617.2, SEM = 17.31) (t(18) = 7.60, p < .001). This indicates that participants used the learned associations between the hiragana cues and the relevant colour to boost search for the target. Due to a technical problem, manual response data was not recorded for one subject in Experiment 2. Here, the learned predictiveness of the hiragana cues from Experiment 1 was extinguished, and now the cues in Experiment 2 never signalled the location of the search target -therefore they cued a distracter. In keeping with an inadvertent bias of attention driven by memory drive here by learned predictiveness of the cues, the now search irrelevant hiragana cues led to slower search responses in this invalid condition (mean = 622.4, SEM = 16.13) relative to the neutral baseline (mean = 606.5, SEM = 15.15) (t(17) = 2.81, p < .012).

### 3.2 fMRI Univariate Results

First, a general linear model was fit to the on-task data to find out the brain areas that showed increased acvitity on cued vs uncued trials, both for Experiment 1 when the hiragana cues were predictive and also in Experiment 2 when the hiragana cues were distracters.

During the on-task periods of Experiment 1 (when the hiragana cues were predictive of the search target) clusters of activity (Z>.3, whole-brain corrected) were found with peaks in MNI −24 8 52, Z=6.99 (left superior frontal gyrus -SFG- and PFC); −44 −50 46, Z= 7.95 (left angular gyrus IPL); 4 −80 −20, Z=5.28 (Cerebellum); 30 −58 58, Z=7.03 (right SPL); and −50 −58 −16, Z=7 (left inferior temporal gyrus).

During the on-task periods of Experiment 2 (when the cues turned into behavioural distracters), whole-brain corrected clusters of activity peaks during the task were found in −2 18 60, Z=6.54 (pre-SMA, left SFG); −48 −52 54, Z=5.11(left angular gyrus, IPL), 62 −46 38, Z=6.84, (right supramarginal gyrus), and −48 8 40, Z=5.03 (middle frontal gyrus).

### 3.3 Pattern Similarity Results

Then neural pattern similarity of task and introspection periods was quantified. A mask was obtained for the whole-brain contrast valid > neutral during on-task blocks involving the regions noted above. Using this mask, voxelwise paratemeter estimates were extracted from the introspection data and from the on-task data, using the unthresholded group-averaged contrast of parameter estimates (cued>neutral). We found that voxel-by-voxel parameters estimates during introspection periods were significantly correlated to the on-task responses (Pearson’s product-moment correlation = 0.753; t(10335) = 116.35, p < .0001). Then, a mask was derived for each of the FSL FEAT output clusters of task-related activity (i.e. valid > neutral) and using the same approach as above, parameter estimates of voxel wise activity were extracted both for the on-task and the introspection datasets to compute the degree of dis(similarity). Figure 2A illustrates these results. Pattern similarity between task and introspection was high in the left inferior temporal gyrus (Pearson=0.691; t(481) =21.004, p < .0001), left superior frontal and PFC (Pearson = 0.784, t(4871) = 88.146, p < .00001, and in the cerebellum (Pearson = 0.768, t(781) = 33.617, p < .00001). Notably, pattern similarity was lower in the right superior posterior parietal lobe (Pearson = 0.156; t(609) = 3.9063, p < .001).

The same pattern of results was replicated in Experiment 2 when the hiragana cues became behavioural distracters (i.e. invalid for search) (see Figure 2B). Using a mask of the whole-brain invalid > neutral contrast from the search data we found that there was a significant correlation between the responses of the voxels within this mask in the task and introspection blocks (Pearson’s product-moment correlation = 0.659; t(4346) = 57.786, p < .0001). Then, similarity analyses were conduced for each of the clusters identified by FSL FEAT. Figure 2B depicts these results. Pattern similarity was high in the left middle frontal gyrus (Pearson = 0.741, t(462) = 23.734, p < .0001, in the right posterior parietal lobe around the supramarginal gyrus (Pearson = 0.681, t(891) = 27.832, p <.0001), in the left posterior parietal around the angular gyrus (Pearson = 0.792, t(1181) = 44.597, p < .0001), middle superior frontal gyrus (Pearson = 0.391, t(1806) = 18.067, p < .0001). These pattern similary scores are depicted in Figure 2.

The above approach provides a similarity metric of the group-averaged activation maps. Further analyses were performed using a within-subject approach (see Methods) in order to accounted for inter-subject variance. The Pearson correlation between on-task and introspection state was computed separately for each subject. The two approaches produce similar results albeit with a lower correlation in the within-subjects approach. Pattern similarity analyses between task and introspection, considering the total brain area activated by the task was found significant in Exp. 1 t(18) = 4.656, p = 0.00019, mean z(r) = 0.38) and also in Exp. 2 t(18) = 3.105, p = 0.0061, mean z(r) = 0.278). Then, using the same within-subject approach we computed the pattern similarity between introspection and on-task states for the different anatomical clusters. In Experiment 1, pattern similarity was significant in all clusters tested except in the cerebellum (left IT t(18) = 2.634, p = 0.017, mean z(r) = 0.248; left IPL, t(18) = 4.394, p = 0.0003, mean z(r) = 0.444; right SPL t(18) = 2.701, p = 0.015, mean z(r) = 0.324; left PFC t(18) = 3.322, p = 0.004, mean z(r) = 0.334; cerebellum t(18) = 2.006, p = 0.06, mean z(r) = 0.207). Thus, the within-subject similarity approach produced similar results to the group-averaged results, with the main exception of right SPL cluster wherein a more dissimilar pattern was found in the group-averaged results but not in the within-subject similarity approach.

**Figure 2:**
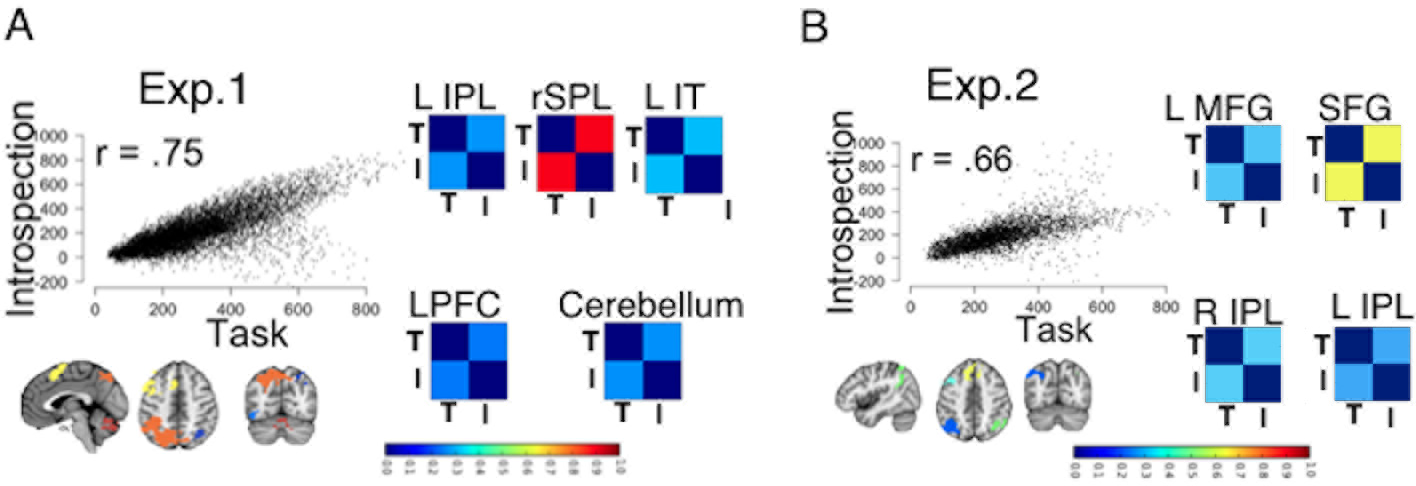
Neural pattern similarity results in Experiment 1 (A) and in Experiment 2 (B). These were conducted using masks from the whole-brain activity clusters in the on-task condition which appear at the bottom left of each panel A and B (see text for further details). Correlation matrices between task (T) and introspection (I) group-averaged contrast of parameter estimates (cued > neutral) are plotted depicting (1 - correlation index; blue is tighter correlation or more similar pattern; red is the reverse).

In Experiment 2, within-subject pattern similarity was in keeping with the group-averaged approach. Similarity between on-task and introspection was significant in all clusters except in the SFG (left IPL, t(18) = 2.359, p = 0.03, mean z(r) = 0.279; right IPL, t(18) = 3.119, p = 0.006, mean z(r) = 0.295; left MFG, t(18) = 2.146, p = 0.046, mean z(r) = 0.217; SFG, t(18) = 2.093, p = 0.051, mean z(r) = 0.203).

### 3.4 Functional Connectivity Results

Because on-task and introspection processes operate on distinct representations (i.e. external/internal) and have distinctive task-based constraints (e.g. perceptual, motor), it is sensible to argue that search and introspection states may be differentiated in how relevant brain areas are functionally related. We therefore turn to the analyses of functional connectivity.

Firstly, we investigated interactions between frontoparietal regions strongly implicated in our fMRI task (see Methods and Figure 3) to explore whether or not a similar connectivity profile occurred during on-task and introspection periods. Finding evidence that the connectivity profile amongst frontoparietal regions can be dissociated across the task and introspection states would rule out accounts that frontoparietal activity during introspection merely reflected domain-general processes (e.g. attentional preparation or working memory demand) triggered by the hiragana cues.

The strength of pairwise functional connectivity for left and right frontoparietal areas was assessed by means of the beta-series correlation method (Rissman, Gazzaley, & D’esposito, 2004). Z-transformed beta-series correlation values within left and within right frontoparietal regions (i.e., IPC, SPC, MFG; see Figure 3A) were averaged and submitted to two separate Hemisphere (left, right) by Context (on-task, introspection) by Cueing condition (cued, neutral) repeated-measures ANOVAs, for both Experiment 1 and Experiment 2 separately.

**Figure 3:**
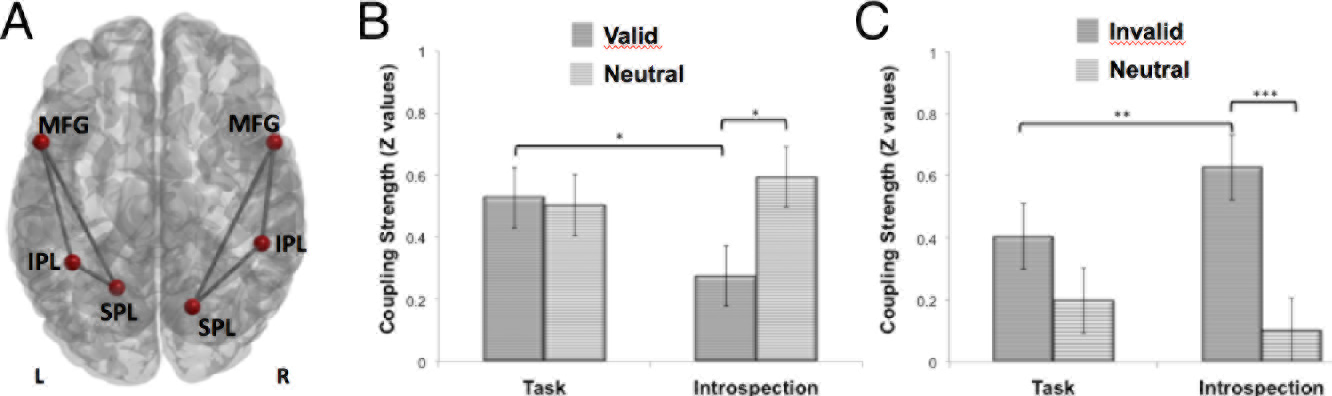
Beta series correlation analysis to assess functional connectivity amogst key frontoparietal nodes activated during on-task periods. (A) A schematic axial view of left and right frontoparietal regions and the pairwise connections among them that were submitted to functional connectivity analysis. All nodes correspond to 5-mm radius spheres centered at the highest local maxima identified from the All-Null contrast, q < .001 FDR voxel-wise corrected for multiple comparisons in left and right IPC, SPC and MFG. Bar graphs show the average coupling strength (i.e., mean Z-transformed values) of the beta-series correlation for the Context * Cueing interactions observed in (B) Experiment 1 and (C) Experiment 2. Asterisks indicate comparisons that showed statistically significant differences in average strength of functional connectivity (* p <. 05; ** p < .01). IPC = inferior parietal cortex; SPC = superior parietal cortex; MFG = middle frontal gyrus).

Results from Experiment 1 showed a main effect of cueing, (F(1, 18) =7.19, p < .05, 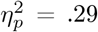), which was qualified by a significant interaction between Context (on-task vs introspection) and cueing, (F(1, 18) = 5.79, p < .05, 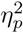 = .24) (see Figure 3B). Post-hoc analysis revealed that this interaction was due to a significant tighter average coupling among frontoparietal areas when participants were validly cued in the task condition relative to the introspection counterpart (p < .05). This difference between on-task and introspection conditions was not observed when participants were presented with neutral cues (p = .24). Also, in the introspection condition stronger average functional connectivity was observed among frontoparietal areas on neutral relative to validly cued trials (p < .05). No significant effect of cueing was observed during on-task periods (p = .70). No main or interactive effects emerged for the factor Hemisphere (Fs(1, 18) = 2.34, ps >.14, 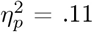), suggesting that the pattern of results was consistent across hemispheres.

Results from Experiment 2 also showed a main effect of Cueing (F(1, 18) =12.5, p < .01, 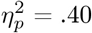), qualified by a significant interaction between Context and Cueing, (F(1, 18) = 4.35, p = .05, 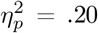) (see Figure 3C). Post-hoc analysis revealed that this interaction was due to a stronger average coupling among frontoparietal areas when participants were cued in the introspection condition compared to when they were cued during the on-task condition (p < .01). This was not observed when participants were presented with neutral cues (p = .46). Also, during introspection there was a tighter average functional connectivity when participants were presented with an invalid cue compared to when they were presented with neutral cues (p < .01), a reversal of the effect relative to Experiment 1 which may be accounted for by the the reversal of the validity or relevance of the cues from Experiment 1 (valid) to Experiment 2 (invalid). No significant differences as a function of Cueing were observed during on-task periods (p = .11). No main or interactive effects emerged for the factor Hemisphere (Fs(1, 18) = .99, ps > .33, 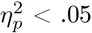).

Lastly, functional connectivity analyses were conducted to test the hypothesis that the integration between the defaul mode network (DMN) and task ‘positive’ control networks is modulated during introspection of our recently past episodes of attentional control. This was done by assessing the temporal correlation between the time courses of each individual’s DMN and the time courses of several brain networks identified by Smith and colleagues (Smith et al., 2009) (see Methods), including the following: visual, auditory, a cerebellum network, sensorimotor, executive control network, left and right frontoparietal networks. and two dorsal attention networks.

The results show that the temporal correlation between the DMN, visual, right frontoparietal and dorsal attention networks increases during introspection relative to the on-task state. This was the case both in Experiment 1 and in Experiment 2. In Experiment 1, the change between DMN and each of the other networks was significant, including visual networks (Vis1: t=3.114, p<.0059; Vis3: =2.680, p<.0153, but not with Vis2: t=1.366, p>0.18), the righ frontoparietal network (t=3.797, p<.0013), DAN1 (t=3.400, p<.0031) and DAN2(t=4.200,p<.0005). DMN also increased its coupling with the sensorimotor network during introspection (t=3.075, p<.0065).

The results are depicted in Figure4, which also highlights the tests that survive multiple comparison correction (Benjamini & Hochberg, 1995).

In Experiment 2, the change between DMN and each of the other networks was significant, including visual networks (Vis1: t = 2.491, p < .023; Vis3: = 3.173, p < .005, but not with Vis2: t = 0.693, p > 0.4), the right frontoparietal network (t = 2.705, p < .014), DAN1 (t = 3.967, p < .0009) and DAN2 (t = 2.667,p < .016).

## 4 Discussion

The main findings can be summarised as follows. Neural responses during introspection of one’s own recently past episodes of attentional performance resembled the pattern of neural responses during the actual behavioural performance. This occurred in a rather distributed set of anatomical locations involving both inferior and superior parietal and superior frontal and prefrontal substrates. Neural pattern similarity between search and introspection states was found in two different task contexts in which the relevance of the attentional cues for behaviour was manipulated, namely, both when the cues served a guiding role in behaviour and when the predictiveness of the cues was extinguished and turned to be associated with the colour of a search distracter rather than the target. This is reminiscent of the cue encoding/retrieval re-activation seen in recognition tests wherein the pattern of cortical activity observed during direct experience with a sensory object is at least partially reinstated when the object is later retrieved or remembered (Danker & Anderson, 2010; Kuhl & Chun, 2014; Nyberg, Habib, McIntosh, & Tulving, 2000; Polyn, Natu, Cohen, & Norman, 2005; Johnson, McDuff, Rugg, & Norman, 2009). Notably, however, the introspection-related demand in this paradigm is distinct from a basic memory requirement (e.g. paired associate learning or delayed recall of items previously encoded), because participants were required to reinstate in their mind the prior experiences during task performance as well the states of cognitive control, and not merely the content-specific task events (e.g. the associations between the hiragana cue and colour of the target).

**Figure 4:**
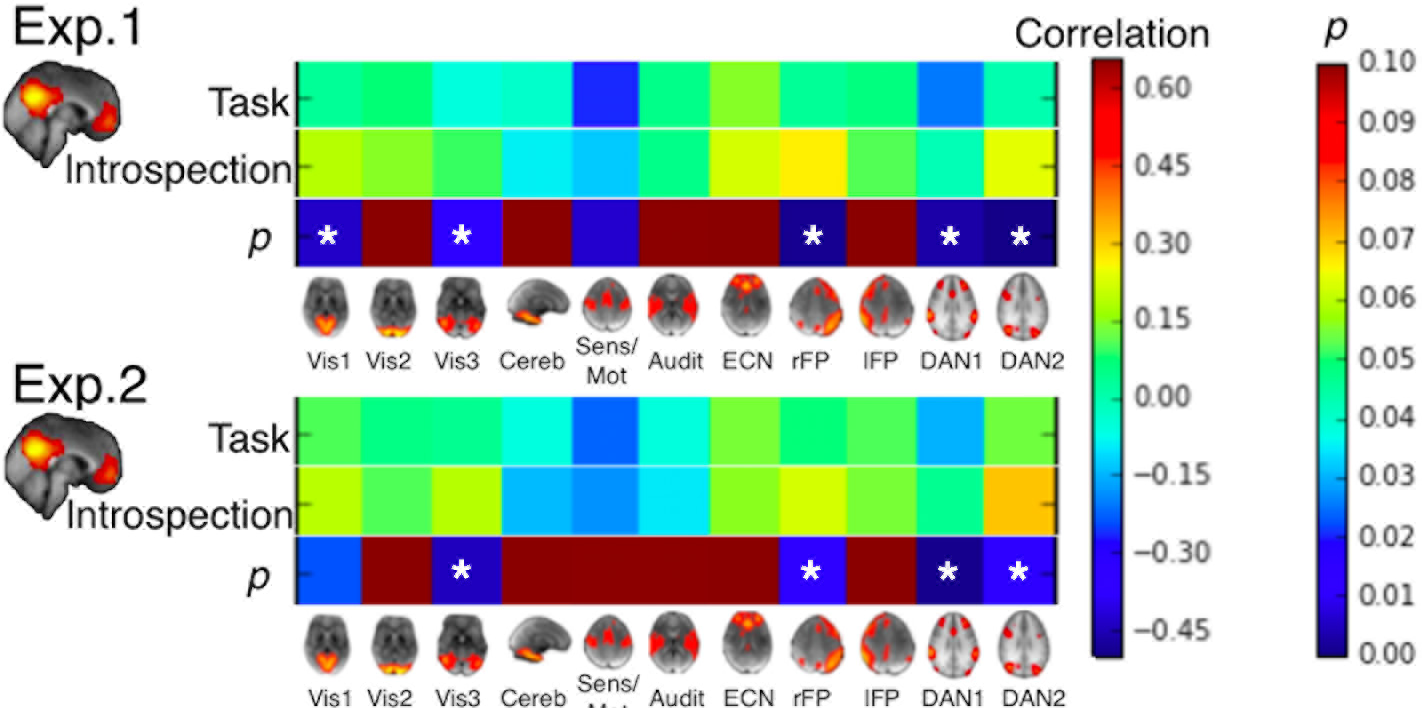
Temporal correlations between the Default mode network (DMN) and several large-scale networks, including three visual (Vis) networks (medial, occipital pole and lateral), cerebellum network, auditory (Audit), sensorimotor (Sensi/Mot), executive control network (ECN), left and right frontoparietal (lFP, rFP), dorsal attention networks (DAN1 and DAN2). The coupling between the DMN and the rest of the networks is illustrated in the above correlation matrix which represents the Pearson correlation between the network time courses, in both the on-task and the introspection periods (* illustrate results corrected for multiple comparisons using FDR).

The pattern of neural activity during the introspection states is also unlikely to merely reflect domain-general processes such as working memory or preparatory attentional control (e.g. similar to those elicited by the hiragana cues during on-task states). These processes would include lingering expectations about the properties of the behavioural target (Corbetta & Shulman, 2002; Hopfinger, Buonocore, & Mangun, 2000; Kastner, Pinsk, De Weerd, Desimone, & Ungerleider, 1999) or the preparation of a critical response (Connolly, Goodale, Menon, & Munoz, 2002; Astafiev et al., 2003), which are known to be mediated by dorsal frontoparietal networks (e.g. intraparietal sulcus and the superior parietal areas, superior frontal gyrus). Several aspects of the results go against this possibility. First, there was a high similarity between the pattern of responses during task and introspection periods in most brain regions during both Experiment 1 and 2. The whole-brain response profile arguably reflected both processes triggered by the cue and by the search display. Second, notably, neural pattern similarity during introspection and on-task periods was high in inferior parietal and temporal regions which are typically involved in target detection rather than preparatory control. Third, in both Experiment 1 and 2, the results showed that the functional connectivity amongst frontoparietal control areas is distinct between introspection and on-task states and that the functional integration of these frontoparietal areas during introspection is sensitive to the behavioural relevance of the cues during the on-task phases. This reversal of the cueing effect seen here relative to Experiment 1 may be accounted for the reversal of the validity or task relevance of the cues from Experiment 1 to Experiment 2. Note that in Experiment 1 the cues were valid hence predicting the target. However, in Experiment 2 this was no longer the case and the cues were associated with a search distracter. Crucially, if the neural correlates of introspection reflected the operation of a similar mechanism to that engaged during on-task states then a similar connectivity profile in this frontoparietal control network would have been expected, but clearly this was not the case.

Together these results indicate that neural pattern similarity between introspection and on-task states did not merely reflect domain-general processes driven by the cue such as working memory or attentional preparation, which could have led to a similar response profile during the on-task state. Rather we suggest that the pattern of results reflects participants’ introspection and re-experience of the cognitive states during task performance. Notably introspection periods were blocked allowing to separate neural activity from the on-task periods. Also, because introspection was blocked, it is unlikely that participants developed expectations of a forthcoming stimulus or action following the presentation of the hiragana cue during introspection blocks. The interpretation that the response profile in frontoparietal areas during introspection reflected participants’ experience and cognitive states during task performance is also consistent with the subjective reports provided by the participants. These indicated that during introspection participants were re-experiencing the task events, reporting that they were thinking about the color association and how this was used to spot the search target and respond to it.

These findings are in keeping with a network memory approach (Fuster, 1997, 2009), according to which memory is a general property of all neural systems reflected in global patterns of brain activity and connectivity for a given experience and behaviour. In this vein, patterns of brain activity during on-task attentional control can be reinstated during introspection of one’s recent behaviour. It is tempting to suggest that large-scale patterns of brain activity engaged during introspection can encode memories of our recent experiences of cognitive control.

The present results are relevant to further understand how the human brain builds and maintains an internal model of one’s own cognition and behaviour. We found that relative to on-task states, during introspection the DMN increases its coupling with frontoparietal control networks, the DAN, and domain-specific regions (i.e. visual cortex). This finding supports the view that functional integration, and segregation, of the DMN and on-task networks is reliant on an internal, self-related focus of experience. In a similar vein, on-task goal-directed behaviour (i.e. during visuo-spatial planning driven by external cues) typically activates frontoparietal and then DAN, however the DMN becomes more coupled with these control networks when participants mentally simulate themselves in a context of goal-directed planning (Spreng, Stevens, Chamberlain, Gilmore, & Schacter, 2010; Gerlach, Spreng, Madore, & Schacter, 2014) and when individuals imagine themselves solving problems (Gerlach, Spreng, Gilmore, & Schacter, 2011).

The idea that self-reflection of one’s recent states of control involves DMN substrates ‘reading’ the state of task-based frontoparietal control networks is consonant with theoretical accounts of self-referential processing that contend an important function of DMN may relate to our experience of being the agent of a cognitive process, such as attending or acting upon the world around us in the context of a sensory-motor loop(Legrand, 2007; Christoff et al., 2011). One limitation of the temporal correlation analyses between the DMN task positive networks is that these comprised the entire scan, rather than being time locked to the onsets of the cue. As such, this analysis provides a bird-view of the pattern of brain network interactions during introspection and how this relates to the pattern of brain interacctions that are observed on-task. Additional work with tailored experimental design is needed to specifically assess how the interplay between DMN and task positive networks during introspection is shaped by the actual relevance or attentional validity of the on-task events that are later re-experienced.

The present findings serve as an empirical foundation to test further hypotheses of how an internal model of one’s own cognitive control processes is built in the human brain. This will require testing self-reflection across multiple contexts of cognitive control to evaluate whether and how DMN patterns of activity and connectivity with key frontoparietal control networks represent the content of experience during task performance and/or the actual processes engaged on-task.

## 5 Acknowledgements

D.S. acknowledges support from the Spanish Ministry of Economy and Competitiveness (MINECO), through the ‘Severo Ochoa’ Programme for Centres/Units of Excellence in R&D (SEV-2015-490), and project grant PSI2016-76443-P which is also funded by the Agencia Estatal de Investigacion (AEI) and Fondo Europeo de Desarrollo Regional (FEDER).

